# A genome-wide miRNA screen identifies regulators of tetraploid cell proliferation

**DOI:** 10.1101/266395

**Authors:** Marc A. Vittoria, Elizabeth M. Shenk, Kevin P. O’Rourke, Amanda F. Bolgioni, Sanghee Lim, Victoria Kacprzak, Ryan J. Quinton, Neil J. Ganem

**Affiliations:** Department of Pharmacology & Experimental Therapeutics, Boston University School of Medicine, Boston, MA 02118, USA.; Department of Biomedical Engineering, Boston University, Boston, MA 02118, USA.; Weill Cornell Medicine/Rockefeller University/Sloan Kettering Tri-Institutional MD-PhD Program, New York, New York, USA; Department of Medicine, Division of Hematology and Oncology, Boston University School of Medicine, Boston, MA 02118, USA.

**Keywords:** Hippo, YAP, NF2, miR-24, polyploid

## Abstract

Tetraploid cells, which are most commonly generated by errors in cell division, are genomically unstable and have been shown to promote tumorigenesis. Recent genomic studies have estimated that ∼40% of all solid tumors have undergone a genome-doubling event during their evolution, suggesting a significant role for tetraploidy in driving the development of human cancers. To safeguard against the deleterious effects of tetraploidy, non-transformed cells that fail mitosis and become tetraploid activate both the Hippo and p53 tumor suppressor pathways to restrain further proliferation. Tetraploid cells must therefore overcome these anti-proliferative barriers to ultimately drive tumor development. However, the genetic routes through which spontaneously arising tetraploid cells adapt to regain proliferative capacity remain poorly characterized. Here, we conducted a comprehensive, gain-of-function genome-wide screen to identify miRNAs that are sufficient to promote the proliferation of tetraploid cells. Our screen identified 23 miRNAs whose overexpression significantly promotes tetraploid proliferation. The vast majority of these miRNAs facilitate tetraploid growth by enhancing mitogenic signaling pathways (e.g. miR-191-3p); however, we also identified several miRNAs that impair the p53/p21 pathway (e.g. miR-523-3p), and a single miRNA (miR-24-3p) that potently inactivates the Hippo pathway via downregulation of the tumor suppressor gene *NF2*. Collectively, our data reveal several avenues through which tetraploid cells may regain the proliferative capacity necessary to drive tumorigenesis.

## Introduction

Cytokinesis, the final stage of cell division, is a highly complex process that cleaves a single cell into two daughter cells following segregation of replicated chromosomes. Failures in cytokinesis gives rise to tetraploid cells, which harbor twice the normal chromosomal content. Proliferating tetraploid cells are highly genomically unstable, and rapidly accumulate both numerical and structural chromosomal abnormalities (Davoli & de Lange, 2011; Ganem & Pellman, 2007; Storchova & Kuffer, 2008). Moreover, tetraploid cells, by virtue of their doubled genomes, are able to both buffer deleterious mutations as they arise and rapidly acquire beneficial mutations that promote adaptation to stressful environments (Dewhurst *et al.*, 2014; Lim & Ganem, 2014; Selmecki *et al.*, 2015; Storchova *et al.*, 2006; Storchova & Kuffer, 2008). Consequently, multiple models have validated that genomically unstable tetraploid cells can promote tumorigenesis, and current estimates indicate ∼40% of human solid tumors pass through a tetraploid intermediate at some stage during their evolution (Davoli & de Lange, 2012; Dewhurst *et al.*, 2014; Fujiwara *et al.*, 2005; Zack *et al.*, 2013).

To combat the potentially oncogenic effects of tetraploidy, non-transformed cells have evolved mechanisms to limit the proliferation of tetraploid cells (Carter, 1967; Ganem *et al.*, 2007; Kuffer *et al.*, 2013). First, it is now known that tetraploid cells activate two core kinases of the Hippo tumor suppressor pathway, LATS1 and LATS2 (Ganem *et al.*, 2014; McKinley & Cheeseman, 2017). Activated LATS1/2 kinases phosphorylate the pro-growth transcriptional co-activators YAP and TAZ, ultimately leading to their cytoplasmic retention and subsequent degradation (Ganem *et al.*, 2014; Moroishi, Hansen, *et al.*, 2015; Pan, 2007). Second, tetraploid cells promote the stabilization of p53 through caspase-2 and LATS2-dependent mechanisms that inhibit the activity of the p53 E3 ubiquitin ligase MDM2 (Aylon *et al.*, 2006; Fava *et al.*, 2017). Stabilization of p53 leads to a corresponding increase in the expression of one of its core target genes, *CDKN1A*, which encodes the CDK inhibitor p21. Consequently, tetraploid cells arrest in G_1_ phase of the cell cycle, after which they commonly senesce (Andreassen *et al.*, 2001; Ganem *et al.*, 2014; Ganem & Pellman, 2007; Panopoulos *et al.*, 2014).

To facilitate tumorigenesis, tetraploid cells must overcome these barriers to cell proliferation (Ganem & Pellman, 2007). There are three broad routes through which tetraploid cells are known to regain proliferative capacity. First, tetraploid cells can proliferate if they have a nonfunctional or defective p53/p21 pathway (Andreassen *et al.*, 2001; Carter, 1967; Ganem *et al.*, 2014; Krzywicka-Racka & Sluder, 2011; Panopoulos *et al.*, 2014). Indeed, analysis of near-tetraploid tumors by the Cancer Genome Atlas (TCGA) reveals a significant enrichment in *TP53* inactivating mutations, demonstrating the importance of nonfunctional p53 in the tolerance of tetraploidy (Crockford *et al.*, 2017).

Second, tetraploid cells can regain proliferative capacity through functional impairment of the Hippo tumor suppressor pathway and a corresponding restoration of YAP activity (Ganem *et al.*, 2014). Analysis of the Cancer Cell Line Encyclopedia (CCLE) has revealed that both YAP overexpression and LATS1/2 deletion are highly enriched in near-tetraploid compared to near-diploid cancer cell lines (Ganem *et al.*, 2014). Importantly, active YAP is sufficient to promote tetraploid cell proliferation even in the presence of a functional p53 pathway (Fava *et al.*, 2017; Ganem *et al.*, 2014).

Finally, tetraploid cells can regain proliferative capability by promoting cyclin D accumulation through hyperactivation of mitogenic signaling, or by amplification of the gene encoding for cyclin D itself (Crockford *et al.*, 2017; Ganem *et al.*, 2014; Potapova *et al.*, 2016). When in excess, cyclin D1 has been shown to inhibit p21 through non-catalytic sequestration, resulting in the formation of cyclin D1/p21 complexes that negate p21-mediated growth inhibition and permit tetraploid proliferation (Crockford *et al.*, 2017).

While functional impairment of the p53/p21 pathway, re-activation of YAP, or increased mitogenic signaling are sufficient to initiate tetraploid proliferation, it remains unclear how cells initially achieve these growth-promoting adaptations. We hypothesized that overexpression of individual microRNAs (miRNAs) may be one route through which tetraploid cells subtly disrupt these pathways to initially escape G_1_ cell cycle arrest. miRNAs, which are small non-coding RNAs, function as critical regulators of gene expression by repressing the translation of complementary messenger RNAs (mRNAs). The complementarity region of miRNAs, known as the seed sequence, is ∼6-8 nucleotides in length and may target multiple sites within the same mRNA and/or simultaneously modulate the translation of hundreds of mRNAs (Bartel, 2009). In this fashion, miRNAs can control the regulation of multiple signaling pathways at once. Indeed, deregulation of miRNA expression is a common feature of human cancers, with overexpression observed in a variety of tumors (Jansson & Lund, 2012).

In this study, we conducted a genome-wide gain-of-function screen to comprehensively identify individual miRNAs that are sufficient to drive tetraploid proliferation when overexpressed. The results of this screen, detailed herein, establish a list of putative oncogenic miRNAs (oncomiRs), many of which have been implicated in driving tumorigenesis. Importantly, we mechanistically define the pathways these miRNAs modulate to overcome tetraploidy-induced G_1_ arrest.

## Results and Discussion

### miRNA Screen to Identify Regulators of Tetraploid Cell Proliferation

We performed a genome-wide gain-of-function screen to identify individual miRNAs that promote the proliferation of G_1_-arrested tetraploid cells. For this screen, we used the diploid, non-transformed retinal pigmented epithelial (RPE-1) cell line as they contain an intact p53 pathway and undergo a durable G_1_ arrest following cytokinesis failure (Ganem *et al.*, 2014). RPE-1 cells were engineered to stably express the fluorescence ubiquitination cell cycle indicator (FUCCI) system. The FUCCI reporter system distinguishes cell cycle position based on a fluorescence readout, where G_1_ cells exhibit red fluorescence (due to expression of a fragment of Cdt1 fused to m-Cherry) and S/G_2_/M cells exhibit green fluorescence (due to expression of a fragment of Geminin fused to Azami-Green) (Sakaue-Sawano *et al.*, 2008).

To generate G_1_-arrested tetraploid cells for the screen, diploid RPE-1 FUCCI cells were treated with 4 μM dihydrocytochalasin B (DCB), an inhibitor of actin polymerization, for 16 hrs to prevent cytokinesis (Ganem *et al.*, 2014). This treatment induces tetraploidy in ∼60% of cells. Fluorescence-activated cell sorting (FACS) was then used to purify tetraploid cells from diploid cells, as previously described (Shenk & Ganem, 2016). In brief, cells exhibiting both 4N DNA content and red fluorescence, indicative of G_1_ tetraploids, were collected (Figure 1A). This methodology results in a highly enriched (>95% pure) tetraploid population (Ganem *et al.*, 2014; Sakaue-Sawano *et al.*, 2008).

**Figure 1:**
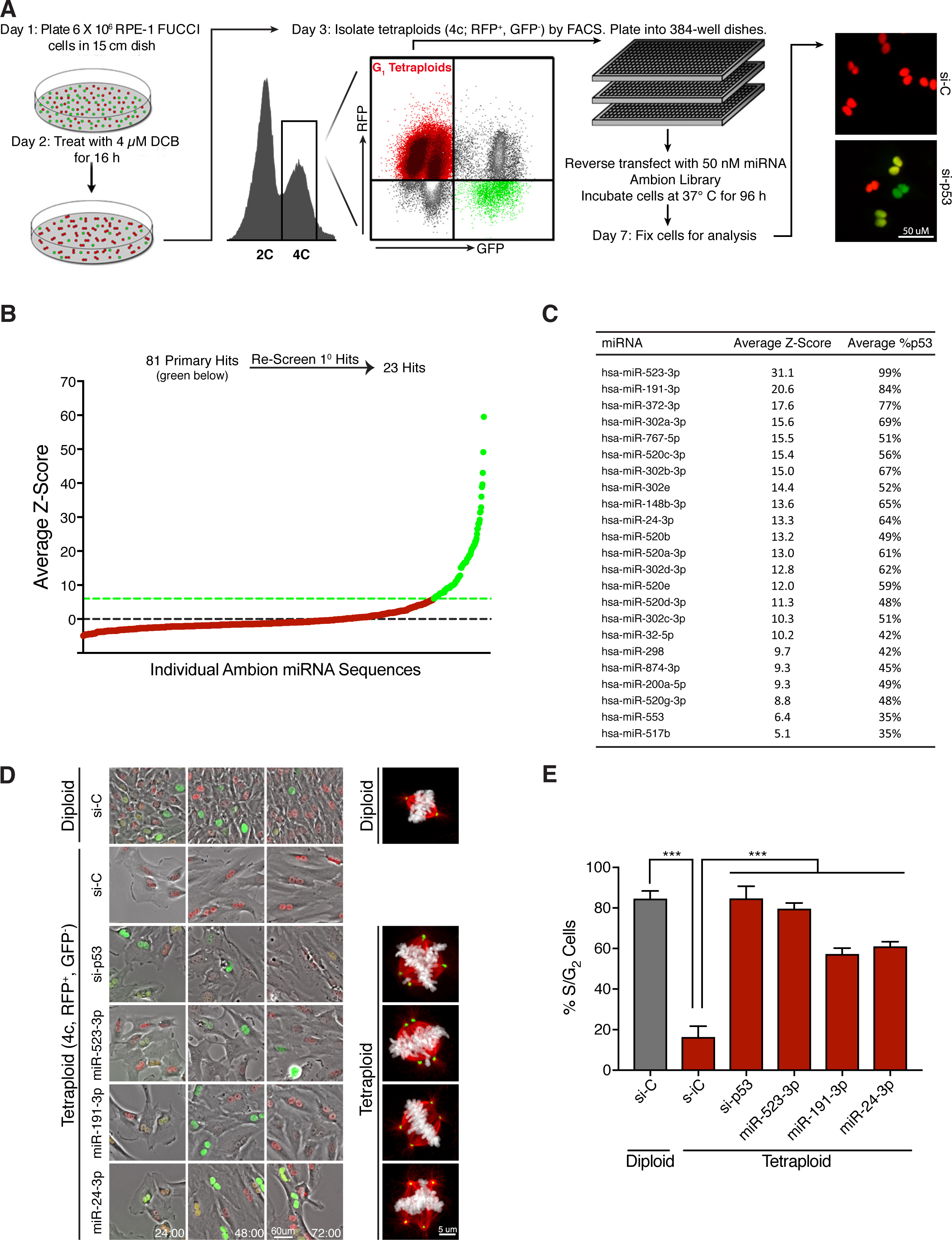
Genome-wide screen to identify miRNAs that promote tetraploid cell proliferation. **(A)** Protocol for a genome-wide miRNA screen to identify overexpressed miRNAs that promote tetraploid cell proliferation. Adapted from Ganem *et al.*, Cell 2014. **(B)** Average Z-scores for the ∼800 screened individual miRNAs in the primary screen. The primary screen yielded ∼80 miRNAs hits (green) that were subsequently re-screened. **(C)** The 23 miRNA hits following secondary screening ranked by average Z-score. **(D)** Still images from a live-cell imaging experiment with RPE-FUCCI cells. Sorted diploids and tetraploids transfected with the indicated siRNA/miRNA are shown at 24, 48, and 72 hr. Representative images of mitotic cells for each condition are shown on right. Microtubules (red), chromosomes (white), centrosomes (green). **(E)** Quantitation of the percentage of cells that enter S/G_2_ at least once for each condition shown in (D) over 72 hr. Error bars represent mean +/- SEM from three separate experiments (n = 50 cells counted per experiment), *** p ≤ 0.001, two-tailed t test and one-way ANOVA.

Consistent with previous studies, we found that the isolated tetraploid cells exhibited a strong G_1_ cell cycle arrest. Live-cell imaging revealed that >80% of all binucleated tetraploid cells failed to enter S-phase over 72 hours, as indicated by persistent red fluorescence (Figure 1, D and E). By contrast, the vast majority of isolated diploid cells, which were exposed to the same culture conditions, drug treatments, and FACS as tetraploids, showed continued proliferation as evidenced by a change from red-to-green fluorescence indicative of the G_1_-S transition (Figure 1, D and E) (Ganem *et al.*, 2014). As a positive control, siRNA-mediated depletion of p53 was observed to restore proliferation to tetraploid cells (Figure 1D).

To perform the screen, freshly purified G_1_-arrested binucleated tetraploids were seeded in 96-well plates in triplicate and reverse transfected with a library of ∼880 precursor miRNA mimics to emulate overexpression of endogenous miRNA (Figure 1A). Non-targeting miRNA sequences were used as negative controls, while siRNAs targeting *TP53* were used as positive controls. At 96 hrs post miRNA transfection, all plates were fixed and automated image analysis was used to determine the fraction of tetraploid cells per well that had escaped G_1_ arrest and entered S-phase (as judged by a transition from red-to-green fluorescence) (Figure 1A). For the primary screen, miRNA hits were identified as having a Z-score ≥6.0, which was determined from multiple intra-plate negative controls. Using this strict criterion, the primary screen identified 81 miRNAs for which overexpression enabled tetraploid cells to escape G_1_ arrest (Figure 1B). To validate and reproduce these hits, each of the 81 miRNAs were then rescreened under the same protocol and manually analyzed. This secondary screen resulted in the identification of 23 miRNAs whose overexpression gave an average Z-score value of ≥5.0 after both biological replicates, indicating robust promotion of tetraploid cell proliferation (Figure 1C). To complement the Z-score ranking, we further determined how well each miRNA promoted tetraploid proliferation relative to RNAi depletion of p53. Remarkably, multiple miRNAs enhanced tetraploid proliferation nearly as well as p53 loss, including the top two hits, miR-523-3p and miR191-3p, which essentially mirrored the ability of p53 knockdown to promote the proliferation of tetraploid cells.

We performed live-cell imaging to confirm that tetraploid cells overcame G_1_ arrest and entered mitosis following transfection with several of our strongest miRNA hits (Figure 1, D and E). As expected, tetraploid cells that entered mitosis frequently displayed abnormal, multipolar spindles as a consequence of containing supernumerary centrosomes (Figure 1D). Thus, our data demonstrate that overexpression of individual miRNAs is sufficient to promote tetraploid cell proliferation and to promote the abnormal mitotic spindle assembly that leads to subsequent genome instability (Ganem *et al.*, 2009; Silkworth *et al.*, 2009).

### The Majority of miRNA Hits Hyperactivate Mitogenic Signaling or Impair the p53/p21 Pathway

As a first assessment of our miRNA hits, we performed cluster analysis to identify whether any shared seed sequence homology. This analysis revealed that over half of the miRNAs share a recently identified oncomotif seed sequence, AAGUGC (Figure S1A). Previous work has demonstrated that miRNAs with an AAGUGC seed motif act as oncomiRs by targeting multiple tumor suppressors to promote the proliferation of cancer cells (Zhou *et al.*, 2017). Thus, this result validated our screening approach and revealed that many clinically relevant oncomiRs may act, at least in part, by promoting the proliferation of tetraploid cells.

We then sought to identify the specific pathways that are disrupted by overexpression of the strongest miRNA hits. Previous studies have demonstrated that hyperactivation of mitogenic signaling is sufficient to promote tetraploid cell proliferation; therefore, we investigated whether any of our miRNA hits enhanced MAPK or PI3K signaling. To do this, RPE-1 cells transfected with scrambled or endogenous miRNA mimics were serum starved for 24 hrs to turn off mitogenic pathways, and then re-stimulated with 5% fetal bovine serum. Following restimulation, both MAPK and PI3K pathway activation was assessed by quantitating the fraction of phosphorylated ERK to total ERK and phosphorylated AKT to total AKT (Figure S1A). Results from this assay revealed that overexpression of miR-191-3p, our second strongest hit, significantly stimulates both the MAPK and PI3K pathways, and leads to a corresponding increase in cyclin D and cyclin E protein levels (Figures 2, A-B and S1, B-C). Furthermore, we found that miR-191-3p overexpression not only hyperactivates the initial mitogenic respsonse to serum, but also maintains a prolonged activation of MAPK and PI3K signaling over multiple hours (Figure 2A). Consequently, cyclin D levels accumulate to a greater extent in miR-191 expressing cells relative to controls (Figure 2B). Due to its overexpression in 16 different cancer subtypes, miR-191-3p is largely classified as an oncogenic miRNA (Di Leva *et al.*, 2013; Nagpal & Kulshreshtha, 2014). Our data reveal that the strong and enduring increase in mitogenic signaling induced by miR-191 overexpression may underpin its capacity to promote tetraploid cell cycle progression, and offer a mechanistic explanation as to its frequent overexpression in human cancers.

**Figure 2.**
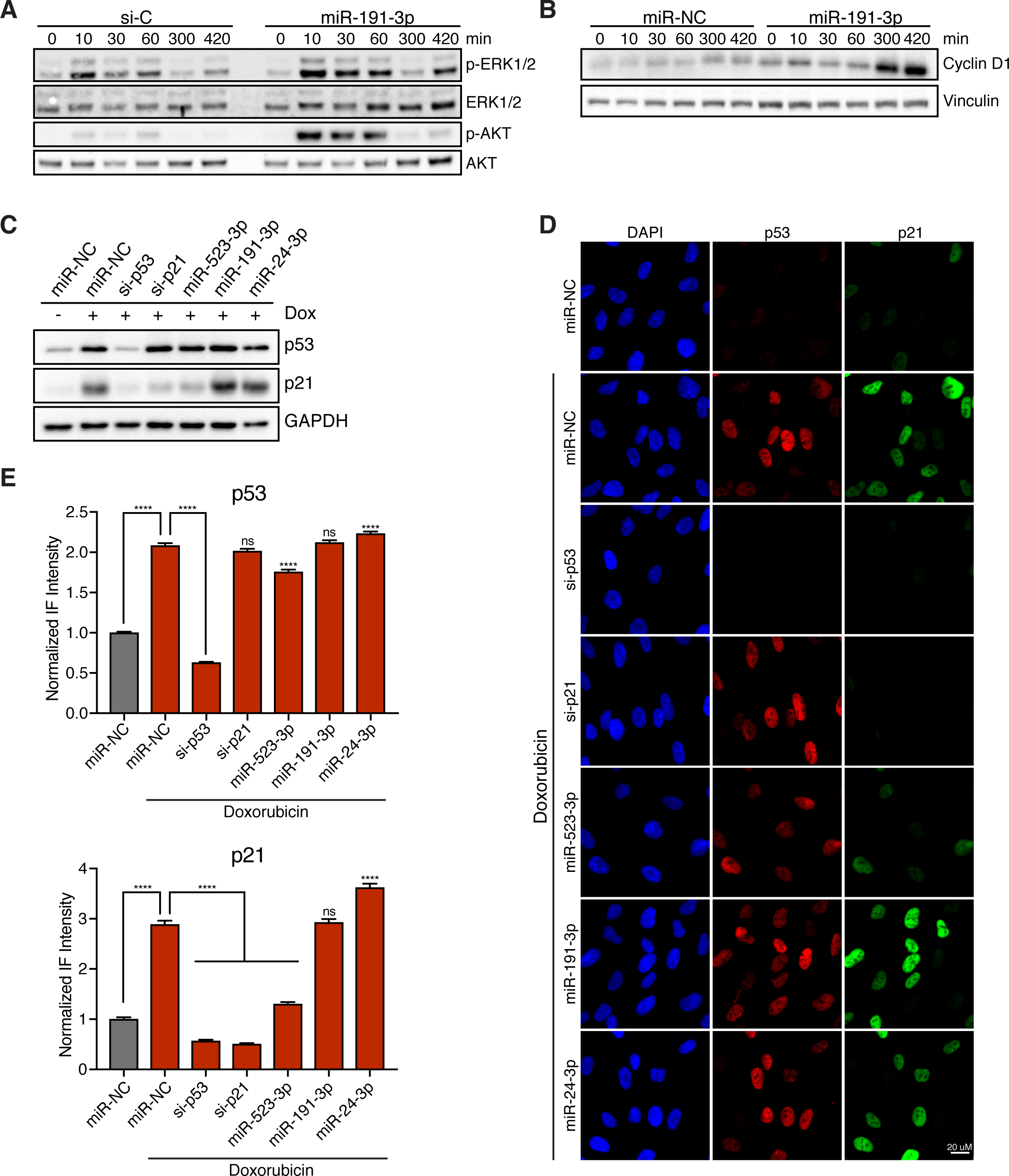
miRNA hits increase mitogenic signaling and/or disable the p53/p21 pathway. **(A)** Representative western blot of RPE-1 cells, transfected with the indicated siRNA/miRNA and serum starved for 24 hr prior to restimulation for the indicated time with 5% fetal bovine serum. **(B)** Representative western of RPE-1 cells, transfected with either non-targeting miRNA control (miR-NC) or miR-191-3p, and serum starved for 24 hours prior to restimulation with 5% fetal bovine serum for the indicated duration. **(C)** Representative western of RPE-1 cells, transfected with the indicated siRNA/miRNA, and treated with or without 100 ng/mL doxorubicin for 4 hr. **(D)** Representative fixed images of RPE-1 cells, transfected with the indicated siRNA/miRNA, and treated with or without 100 ng/mL doxorubicin for 4 hrs prior to fixation. **(E)** Normalized fluorescence intensity quantification of images in (D). Error bars represent mean +/- SEM (n > 650 cells/condition from one experiment), **** p ≤ 0.0001 and ns = non-significant, two-tailed t test and one-way ANOVA.

In addition to hyperactive mitogenic signaling, abrogation of the p53/p21 signaling axis is also known to restore proliferation to tetraploid cells (Figure 1E) (Fujiwara *et al.*, 2005; Ganem *et al.*, 2014; Krzywicka-Racka & Sluder, 2011). Thus, we investigated if overexpression of any of the strongest miRNA hits disrupts p53-induced increase in p21 protein. In normal RPE-1 cells, endogenous levels of p53 and p21 are quite low, making detection of decreases in their protein levels difficult. To address this, control cells or cells transfected with miRNA hits were first treated with 100 ng/ml of the DNA damaging agent doxorubicin for 4 hrs to induce the p53/p21 pathway. In control cells, this treatment strongly activated p53 and elicited a three-fold increase in p21 protein levels (Figure 2, C-E). Using this approach, we identified that our strongest overall hit, miR-523-3p, significantly reduces p21 protein levels (Figures 2, C-E and S1D). This effect is likely to be indirect, as bioinformatics approaches do not strongly predict miR-523-3p to target the 3’ untranslated region of *CDKN1A* mRNA, which we then confirmed with luciferase assays (data not shown). Thus, our data reveal that disruption of normal p53/p21 signaling by overexpression of individual miRNAs is another route through which tetraploid cells can escape cell cycle arrest.

### Overexpression of miR-24-3p Promotes YAP Activation and Tetraploid Cell Proliferation

Functional inactivation of the Hippo tumor suppressor pathway, which leads to YAP activation, is also known to confer proliferative ability to tetraploid cells (Ganem *et al.*, 2014). Therefore, we explored whether overexpression of any miRNA hits is sufficient to increase YAP activity. To do this, we quantitated YAP localization in cells transfected with miRNA hits as an indirect readout of its activity, as it is known that YAP is predominantly cytoplasmic (inactive) upon Hippo pathway activation, but predominantly nuclear (active) upon Hippo pathway inactivation. (Pan, 2007; Yu *et al.*, 2015). Utilizing this assay, we identified two miRNAs (miR-24-3p and miR-553) that significantly increase nuclear YAP localization relative to controls, indicative of Hippo pathway attenuation (Figures 3, A-B and S2A). Interestingly, miR-24-3p overexpression results in more nuclear-localized YAP than even RNAi-mediated depletion of the kinases LATS1/2, which served as the positive control in this assay. This finding was reproduced in HEK293A cells, a cell line frequently employed to study Hippo signaling (Figure S2, B and C). (Meng *et al.*, 2015; Moroishi, Park, *et al.*, 2015). In contrast to miR-523-3p and miR-191-3p, overexpression of miR-24-3p did not abrogate the p53/p21 pathway nor did it activate mitogenic signaling (Figure 2C and S1B).

**Figure 3:**
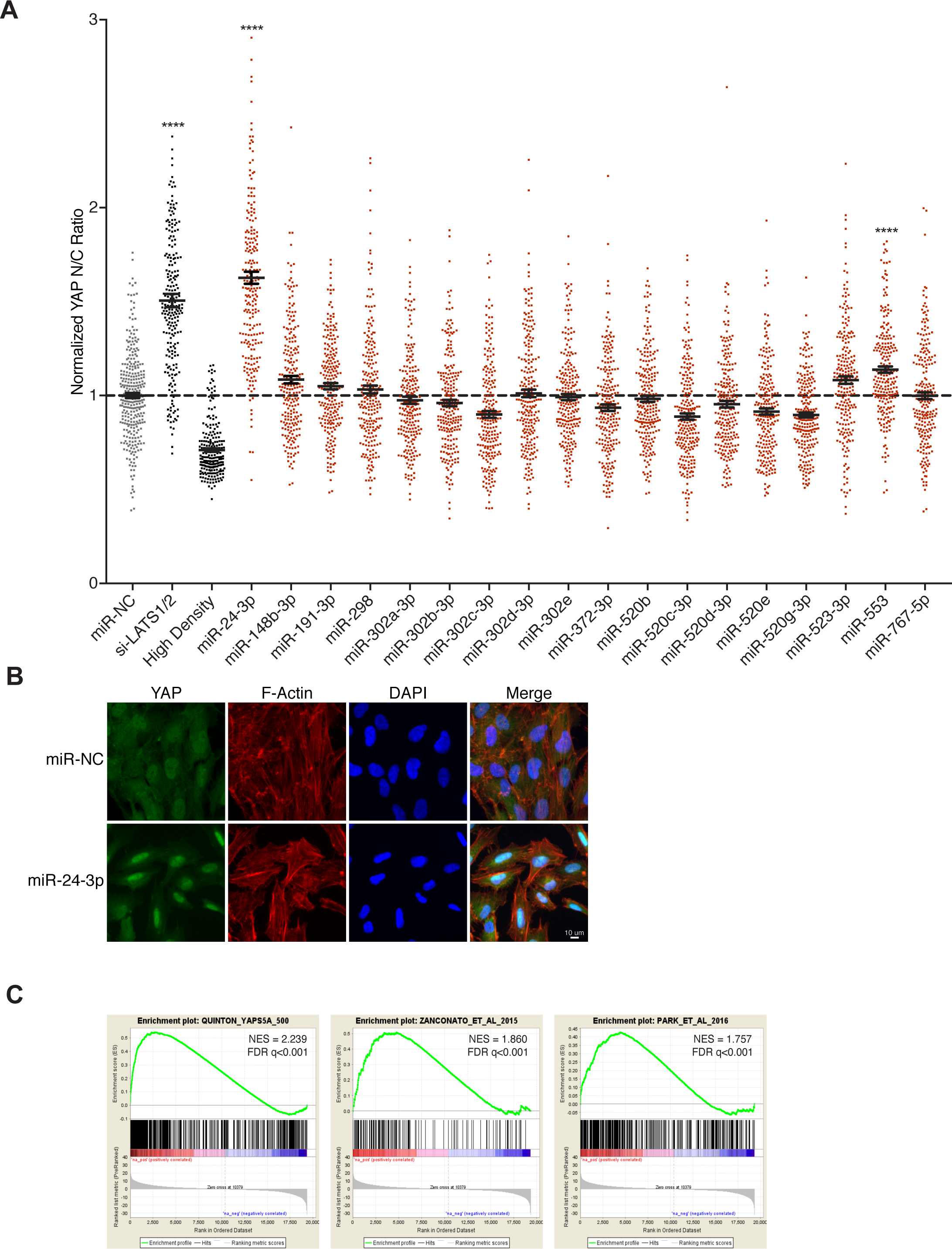
miR-24 overexpression promotes YAP activation. **(A).** Relative localization of YAP in RPE-1 cells following transfection with the indicated siRNA/miRNA. Plot depicts the normalized ratio of YAP immunofluorescence intensity in the nucleus:cytoplasm (N/C) performed in triplicate (n = 3). Error bars represent mean +/- SEM (n >200 cells/condition, **** p ≤ 0.0001, one-way ANOVA). **(B)** Representative fixed images of YAP localization 48 hr following transfection with miR-24-3p or negative control (n > 3) **(C)** Gene-set enrichment analysis of RPE-1 cells transfected with miR-24-3p compared to RPE-1 cells expressing YAPS5A and two YAP dependent gene-sets.

To examine whether the observed increase in nuclear YAP by miR-24-3p overexpression led to a corresponding increase in transcription of YAP-regulated target genes, we performed gene expression analysis. We compared the gene expression profiles of three cell lines: control RPE-1 cells, RPE-1 cells overexpressing miR-24-3p, and RPE-1 cells expressing a constitutively active version of YAP in which five serines phosphorylated by LATS are mutated to alanines (YAPS5A). Gene set enrichment analysis (GSEA) of the resultant expression data confirmed that YAP-target genes, as defined by the RPE-1 YAP-S5A cells and previously published YAP-dependent gene sets, are significantly upregulated in RPE-1 cells expressing miR-24-3p (Figure 3C) (Park *et al.*, 2016; Zanconato *et al.*, 2015). Thus, our data reveal that overexpression of the miRNA miR-24-3p leads to YAP activation and tetraploid cell proliferation (Figure 1D).

### miR-24-3p Overexpression Results in Downregulation of the Tumor Suppressor *NF2*

We aimed to identify potential targets of miR-24-3p to elucidate how it impairs the Hippo pathway to activate YAP. First, we assessed the effect of miR-24-3p overexpression on the levels of the most well characterized Hippo pathway components (e.g. LATS1, LATS2, MST1, MST2; Figure S2D). However, none of the core Hippo kinases were downregulated due to miR-24-3p overexpression. We then focused our attention on our gene expression data, where we found that an upstream activator of the Hippo pathway, the tumor suppressor gene Neurofibromin 2 (*NF2*), was one of the 50 most downregulated genes upon miR-24-3p overexpression (Figure S2E). Known as Merlin in *Drosophila*, NF2 promotes activation of the LATS1/2 kinases, resulting in the phosphorylation and subsequent cytoplasmic sequestration of YAP. Loss of NF2 reduces Hippo pathway activity and promotes active, nuclear YAP (Meng *et al.*, 2016; Moroishi, Park, *et al.*, 2015). This led us to test whether miR-24-3p overexpression activates YAP through downregulation of *NF2*.

Consistent with our expression data, we found that miR-24-3p overexpression results in an over 50% decrease in NF2 protein levels (Figure 4, A and B). To compare relative YAP activation between miR-24-3p overexpression and depletion of NF2, we performed immunofluorescence imaging of YAP localization following treatment with siRNAs targeting NF2. Indeed, RNAi knockdown of NF2 significantly increases nuclear YAP localization; however, the observed effect does not recapitulate the magnitude of increased nuclear YAP localization achieved by miR-24-3p (Figure 4, C and D). These data suggest that miR-24-3p likely targets multiple mRNAs to inactivate the Hippo pathway and promote YAP activity.

**Figure 4:**
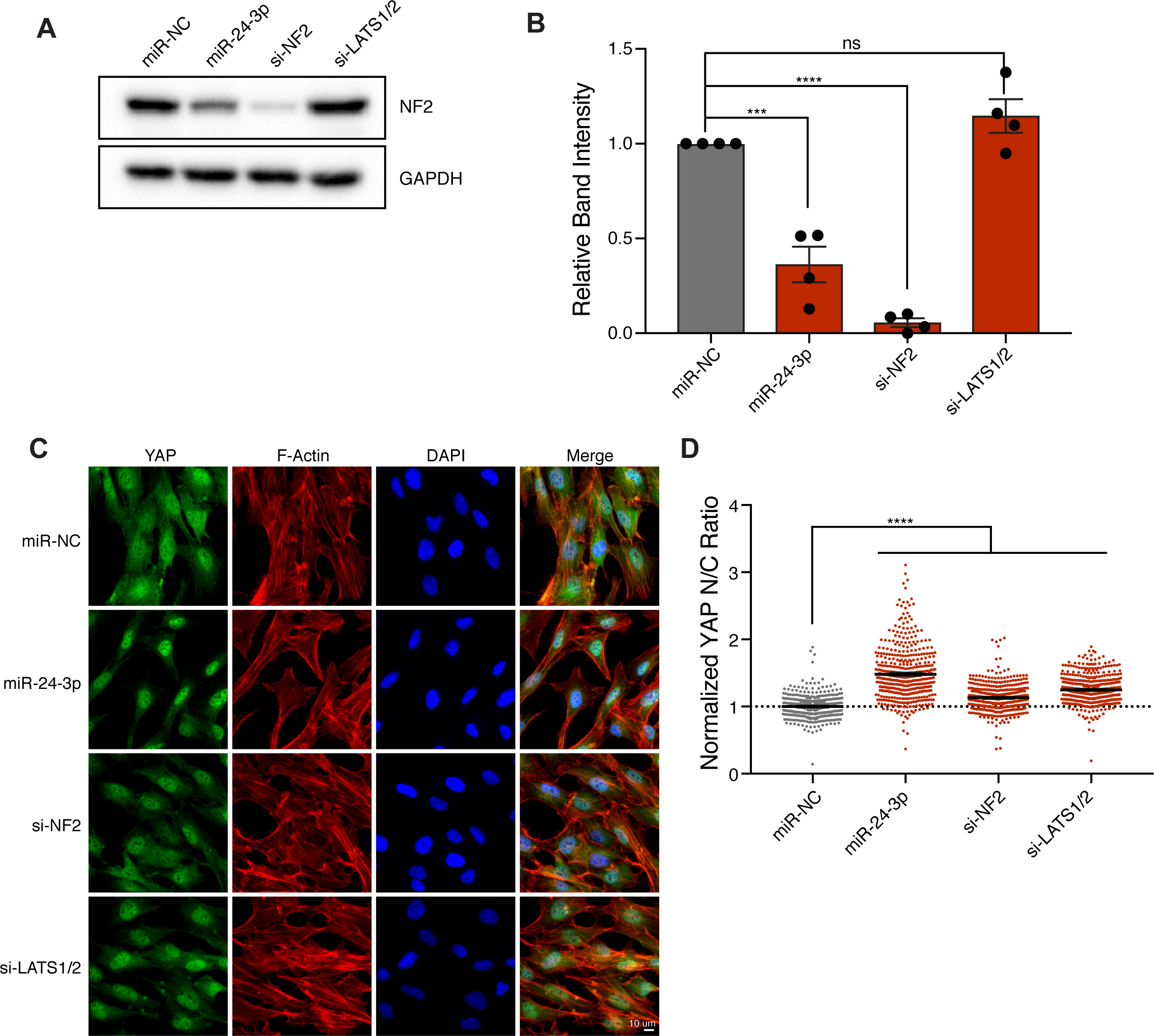
miR-24 activates YAP through down regulation of *NF2*. **(A)** Representative western blot of NF2 protein levels in RPE-1 cells transfected with the indicated siRNA/miRNA after 48 hr. **(B)** Quantification of western blots in (A). Error bars represent mean +/- SEM (n > 3, *** p ≤ 0.001, **** p ≤ 0.0001, ns = non-significant, one-way ANOVA). **(C)** Representative fixed images of YAP localization in RPE-1 cells transfected with the indicated siRNA/miRNA after 48 hr (n > 3). **(D)** Plot depicts the normalized ratio of YAP immunofluorescence intensity in the nucleus:cytoplasm (N/C) from (C). (n > 450 cells/condition, **** p ≤ 0.0001, one-way ANOVA).

Previous work has identified multiple roles for miR-24-3p in carcinogenesis. miR-24 is overexpressed in many cancer subtypes (e.g. breast, hepatic, and Hodgkin lymphoma) and its upregulation promotes cell proliferation through disruption of the cyclin-dependent kinase inhibitors p27^Kip1^ and p16^INK4a^. (Giglio *et al.*, 2013; Hatziapostolou *et al.*, 2011; Kerimis *et al.*, 2017; Liu *et al.*, 2017; Yuan *et al.*, 2017). Interestingly, small RNA sequencing has revealed several miRNAs that are highly overexpressed in patient hepatocellular carcinoma samples relative to cirrhotic nodules, including miR-24-3p with an over 7-fold increase in expression (Sulas *et al.*, 2017). This is relevant since hepatocytes from adult animals are predominantly tetraploid, and YAP activation has been shown to promote their proliferation *in vivo* and encourage the development of HCC (Ganem *et al.*, 2014; Patel *et al.*, 2017; Zhang *et al.*, 2017). As such, our data suggest that miR-24-3p overexpression in tetraploid hepatocytes may promote cell cycle progression through activation of YAP, thus providing one potential explanation for this clinical observation.

### Summary

Here, we identify 23 miRNAs whose expression enables tetraploid cells to escape G_1_ arrest and re-enter the cell cycle. We demonstrate that these miRNAs promote tetraploid cell proliferation by either hyperactivating mitogenic signaling, disrupting the p53/p21 pathway, or functionally impairing the Hippo tumor suppressor pathway to activate YAP. Our data suggest that subtle modulation of miRNA expression levels, even transiently, provides routes through which tetraploid cells can overcome cell cycle arrest and regain proliferative capacity. As proliferating tetraploids are highly genomically unstable, this initial proliferative push may set in motion a subsequent series of mutational events that ultimately lead to tumorigenesis.

## Materials and Methods

### Cell Culture

Telomerase-immortalized RPE-1 cells (ATCC), and all derivative cell lines generated in this study, were grown in phenol red-free DMEM:F12 media containing 10% FBS, 100 IU/ml penicillin, and 100 μg/ml streptomycin. All cells were maintained at 37°C with 5% CO_2_ atmosphere.

### Viral Infections and siRNA Transfections

RPE-1 cells were infected for 12-16 h with virus carrying genes of interest in the presence of 10 μg/ml polybrene, washed, and allowed to recover for 24 hr before selection. All siRNA transfections were performed using 50 nM siRNA with Lipofectamine RNAi MAX according to the manufacturer’s instructions.

### Tetraploid miRNA Screen

**Day 1**: 15 cm dishes were seeded with 6 million exponentially growing RPE-FUCCI cells, such that they were ∼65% confluent the following day. **Day 2**: 4 μM DCB was added to each 15 cm dish for 16 hr. **Day 3**: DCB-treated cells were washed 5 × 5 min, incubated in medium containing 2.5 μg/ml Hoechst dye for 1 hr, trypsinized in 0.05% trypsin, pelleted, resuspended in fresh medium, and FACS sorted. G_1_ diploids (2C DNA content; mCherry^+^, GFP^-^) and G_1_ tetraploids (4C DNA content; mCherry^+^, GFP^-^) were isolated by FACS. Sorted cells were pelleted, re-suspended in fresh medium without antibiotics, and re-plated at a density of 5000 cells per well of a 96-well plate. Cells were then reverse-transfected using Lipofectamine RNAi Max (according to the manufacturer’s specifications). The final concentration of miRNAs per well was 50 nM. Each plate was screened in triplicate, and was internally controlled with multiple p53 siRNA-positive controls and scrambled miRNA negative controls. **Day 4**: All transfected wells were fed with fresh medium containing penicillin/streptomycin. **Day 6**: 100 μM monastrol was added to each well for 12 hr to arrest proliferating GFP^+^ cells in mitosis. **Day 7**: 96 hr following transfection, cells were fixed with 4% paraformaldehyde. Fluorescent images from each well were acquired using a Nikon TE2000-E2 inverted microscope equipped with a cooled CCD camera (Orca ER, Hamamatsu), and Nikon Perfect Focus. An encoded precision stage was used to capture 9 fields of view from each well of the 96-well dish. Subsequently, the total number of proliferating S/G_2_/M cells (based on GFP positivity of the FUCCI system) was calculated as a fraction of the total number of cells (based on nuclear counts using Hoechst) for each well. The Z-score was calculated using the formula 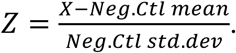 All miRNAs that had a median Z-score of 6.0 were re-tested in a secondary screen, under the same protocol. For the secondary screen, all miRNAs that had a median Z-score above 3.0 were manually examined, and any miRNAs that caused cell death or yellow binucleates (indicative of a failure of the FUCCI degradation system) were removed. The Z-scores from the two screens were then averaged for each miRNA, resulting in the list of hits.

### Immunofluorescence Microscopy

All cells were washed in PBS and fixed in 4% paraformaldehyde for 10 min. Cells were extracted in PBS-0.5% Triton X-100 for 5 min., blocked for 30 min in TBS-BSA (10 mM Tris, pH 7.5, 150 mM NaCl, 5% BSA, 0.2% sodium azide), and incubated with primary antibodies diluted in TBS-BSA for 30-60 min in a humidified chamber. Primary antibodies were visualized using species-specific fluorescent secondary antibodies (Molecular Probes) and DNA was detected with 2.5 µg/ml Hoechst. Immunofluorescence images were collected on a Nikon Ti-E inverted microscope (Nikon Instruments). Immunofluorescence images of RPE-FUCCI cells, including all data collected from the RNAi screen, were captured on a Nikon TE2000-E2 inverted microscope. Confocal immunofluorescence images were collected on a Nikon Ti-E inverted microscope equipped with a C2+ laser scanning confocal head with 405 nm, 488 nm, 561 nm, 640 nm laser lines. Images were analyzed using NIS-Elements software and ImageJ. To assess YAP localization, two small boxes were drawn in individual cells: one in the nucleus, and one in the cytoplasm. The mean fluorescence intensity of YAP was measured in these regions of interest and a nuclear:cytoplasmic ratio was determined. All quantifications of YAP fluorescence localization were completed in a blinded fashion.

### Live-cell Imaging

RPE-FUCCI cells were grown on glass-bottom 12-well tissue culture dishes (Mattek) and imaged on a Nikon TE2000-E2 inverted microscope equipped with the Nikon Perfect Focus system. The microscope was enclosed within a temperature- and CO_2_-controlled environment that maintained an atmosphere of 37°C and 5% humidified CO_2_. Fluorescence and phase contrast images were captured every 10-20 minutes with a 10X 0.5 NA Plan Fluor objective, at multiple points for 2-4 days. All captured images were analyzed using NIS-Elements software.

### Protein Extraction, Immunoprecipitation and Immunoblotting

Cells were washed twice in ice-cold PBS and lysed using ice-cold Laemmli Lysis Buffer containing HALT (dual phosphatase and protease inhibitor, Thermo Fisher). Lysates were sonicated at 20% amplitude for 20 seconds, diluted in 4X Sample Buffer (Boston BioProducts), and resolved using SDS gel electrophoresis. Proteins were then transferred onto PVDF membranes, blocked for 1 h with TBS-0.5% Tween containing 5% skim milk powder, and then probed overnight at 4°C with primary antibodies. Bound antibodies were detected by horseradish peroxidase-linked secondary antibodies and processed with ECL (Amersham) or Clarity ECL (Bio-Rad). Chemiluminescence acquisition was carried out using the Bio-Rad ChemiDox XRS+ system and analyzed using Bio-Rad Image Lab.

### Growth Factor Re-stimulation

Control RPE-1 cells, or RPE-1 cells transfected with miRNAs/siRNAs for 48 h, were serum-starved overnight; they were then stimulated with medium containing 5% serum for the indicated time points, then collected immediately for quantitative western blot analysis.

### Microarray Analysis and GSEA

Total RNA was extracted from exponentially growing control RPE-1 cells (transfected with scrambled siRNA), RPE-1 cells expressing YAP-5SA, and RPE-1 cells transfected with miR-24-3p for 48 hr with the RNeasy kit (Qiagen) and hybridized onto Affymetrix HG-U133_Plus_2 arrays as per the manufacturer’s instructions. Gene set enrichment analysis (GSEA) was performed using the GSEA application from the Broad Institute. Analysis was performed using preranked gene lists with phenotype as permutation type, 1,000 permutations and log_2_ ratio of classes as metric for ranking genes (Mootha *et al.*, 2003; Subramanian *et al.*, 2005). YAP/TAZ dependent gene-sets were curated from the following expression data: GSE66083 and GSE54617 (Park *et al.*, 2016; Zanconato *et al.*, 2015).

### Reagents and Antibodies

The following antibodies were from Cell Signaling Technologies: AKT (#9272), Phospho-AKT (#4051), ERK (#9107), Phospho-ERK (#9101), LATS1 (#3477), LATS2 (#5888 and #13646), p21 (#2947), YAP (#4912), MST1 (#3682), MST2 (#3952), NF2 (#12896), and GAPDH (#2118). Antibodies against p53 (DO-1), p21 (F-5), Cyclin D1 (A-12), Cyclin E (M-20), MARK2 (H-86) and YAP (63.7) were from Santa Cruz. Antibody against beta-actin (AC-74) was from Sigma. Antibodies against Vinculin (ab18058) and AMOTL1 (ab84049) were from Abcam. Doxorubicin and Dihydrocytochalasin B were from Sigma.

### Plasmids

Plasmids encoding cDNA for YAP-S5A (#33093) was obtained from Addgene. YAP-S5A was cloned into the pBABE backbone, making use of PCR-based cloning (In-Fusion HD, Clontech).

### Statistics

Prism 7 was used for all statistical analyses and for the creation of all graphs.

### miRNA and siRNA Sequences

The miRNA library was from Ambion. Accession numbers and sequences for hits are listed below.

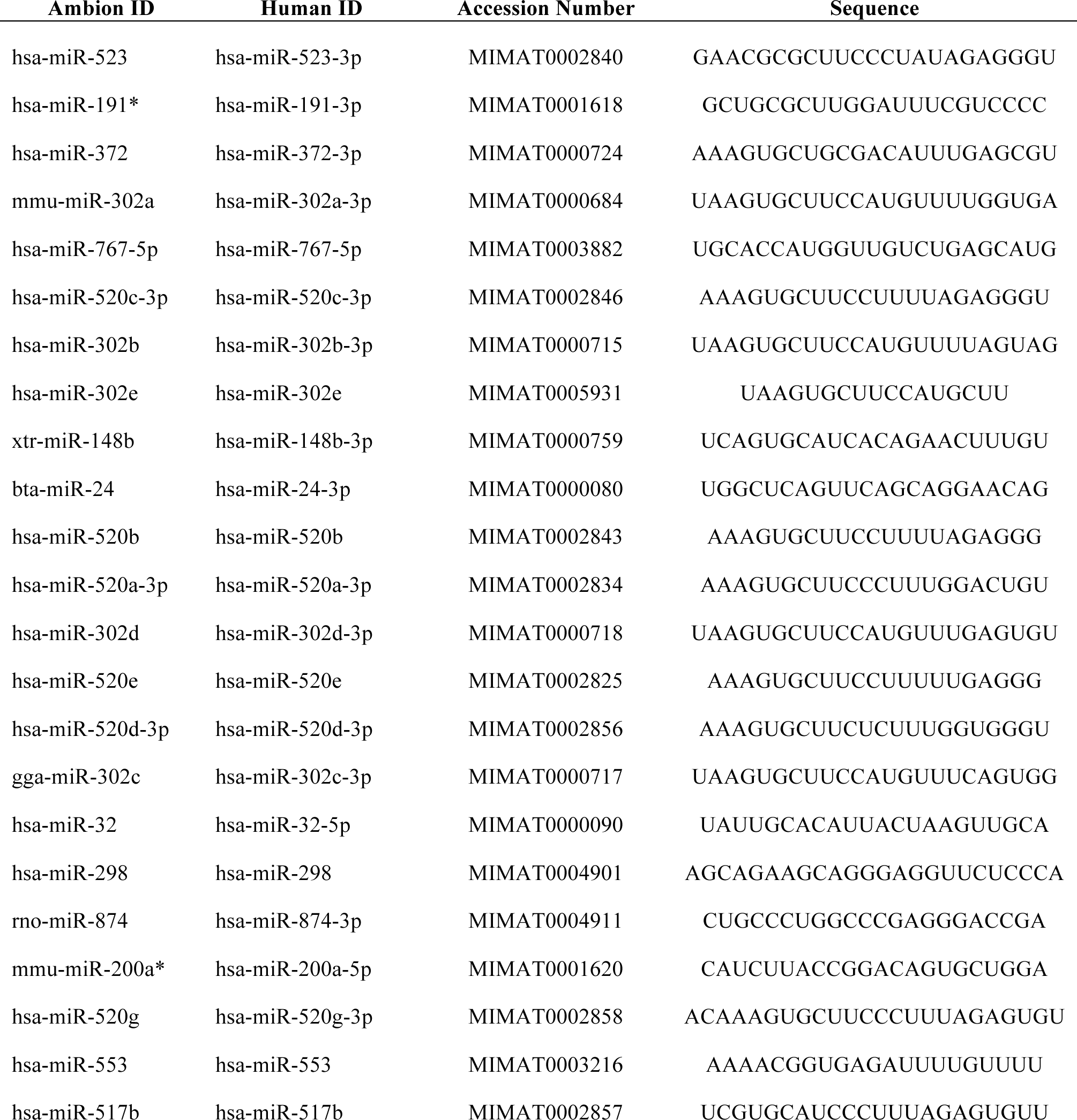

The siRNAs used in this study:

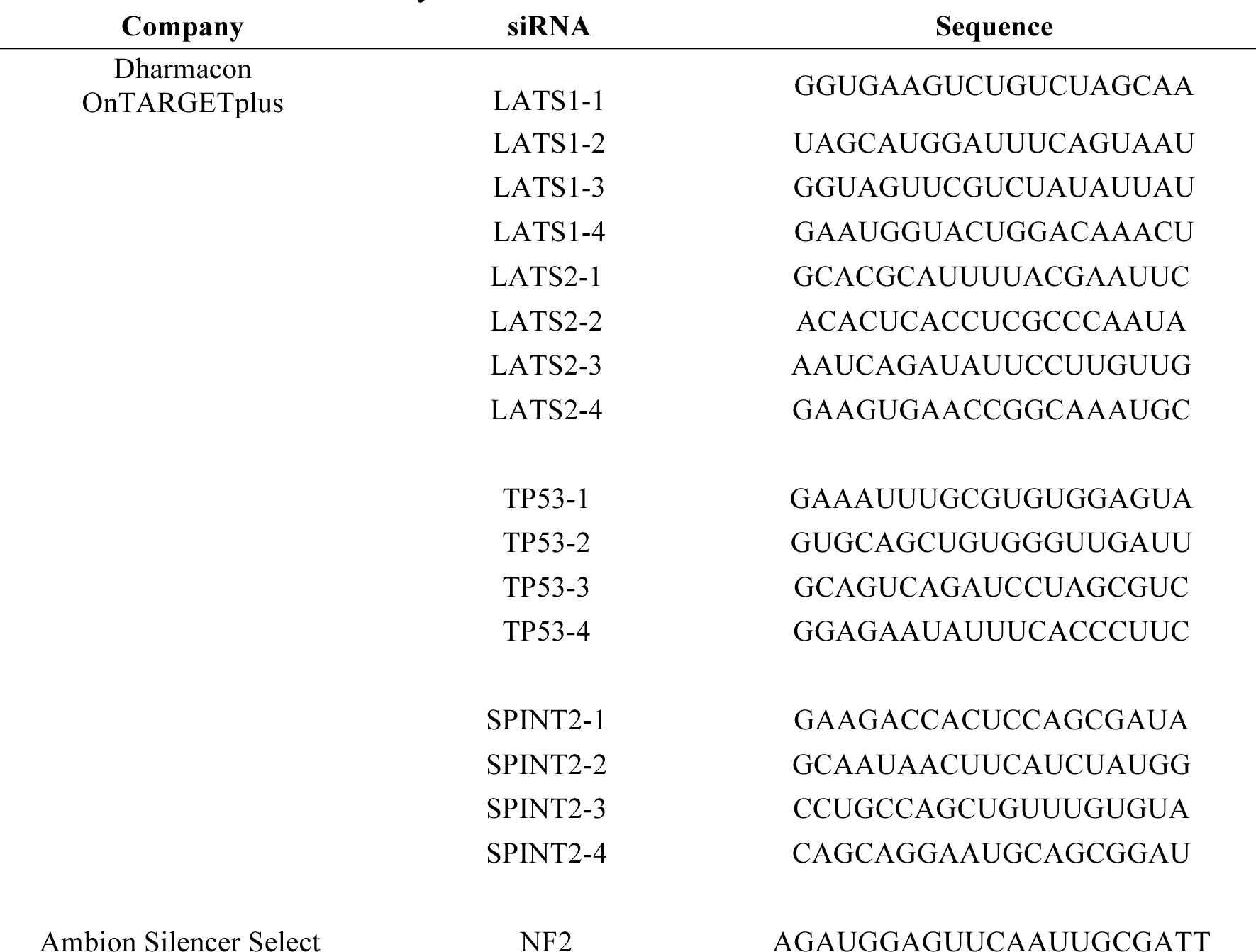

## Acknowledgments

We would like to thank members of the Ganem lab for comments on the manuscript and members of the ICCB screening facility at Harvard Medical School for their expertise in high-throughout screening. M.V., E.S., and A.B. were supported by a training grant from the NIH NIGM (5T32GM008541-20). RQ is supported by the Canadian Institutes of Health Research Doctoral Foreign Study Award. K.O. is supported by an F30 Award from the NIH/NCI (1CA200110-01A1). N.G is a member of the Shamim and Ashraf Dahod Breast Cancer Research Laboratories and is supported by NIH grants CA154531 and GM117150, the Karin Grunebaum Foundation, the Smith Family Awards Program, the Melanoma Research Alliance, and the Searle Scholars Program.

---

**Supplemental Figure 1:**
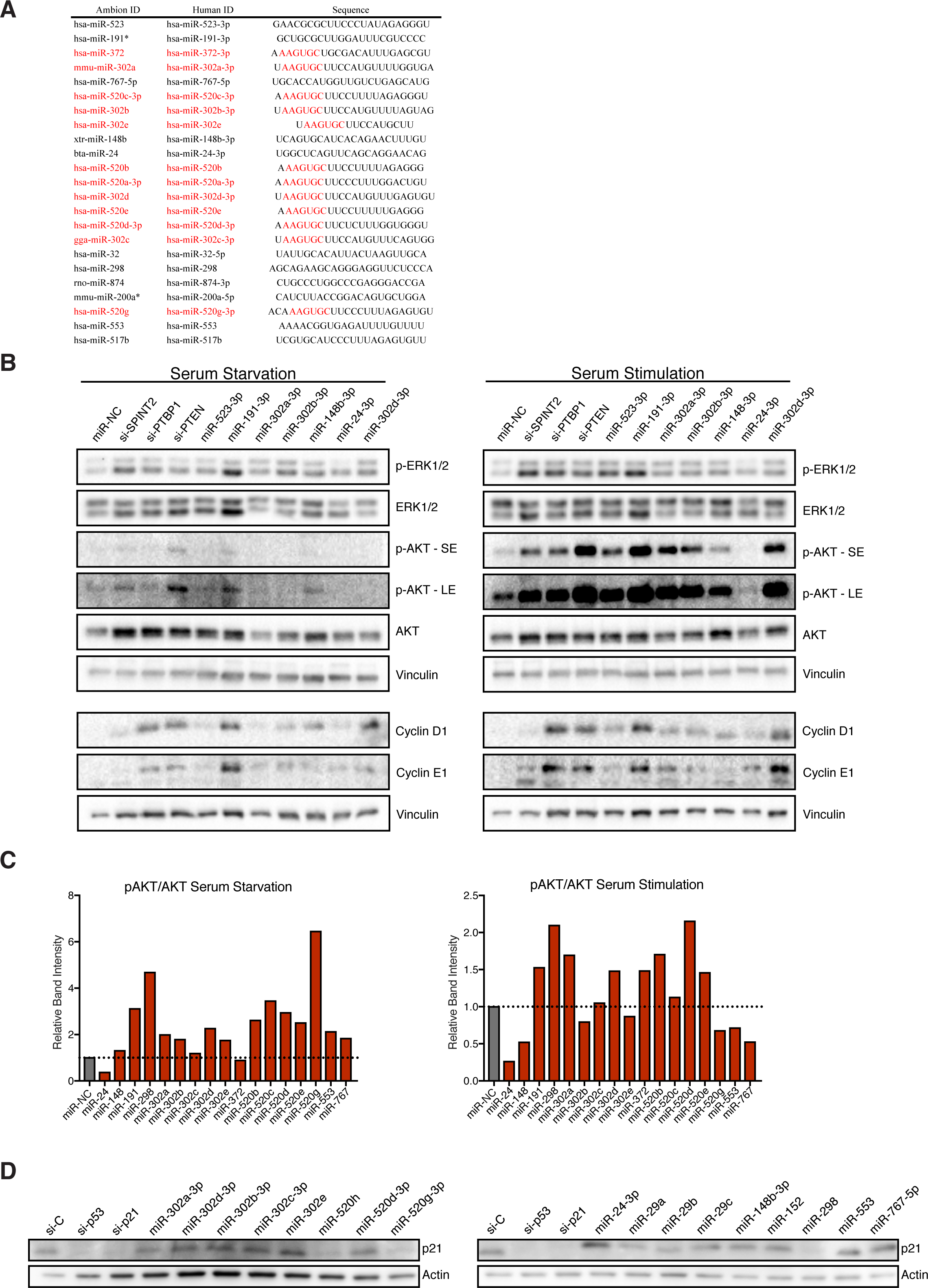
**(A)** Table identifying all miRNA hits with an AAGUGC seed motif. **(B)** Western blot analysis of RPE-1 cells, transfected with the indicated siRNA/miRNA and serum starved for 24 hrs (left) prior to 1 hr restimulation with 5% fetal bovine serum (right). **(C)** Quantification of phosphorylated AKT over total AKT levels in RPE-1 cells, transfected with the indicated siRNA/miRNA and serum starved for 24 hr (left) prior to 1 hr restimulation with 5% fetal bovine serum (right) (n = 1). **(D)** Western blot analysis of RPE-1 cells, transfected with the indicated siRNA/miRNA and after treatment with 100 ng/mL doxorubicin for 4 hrs.

**Supplemental Figure 2:**
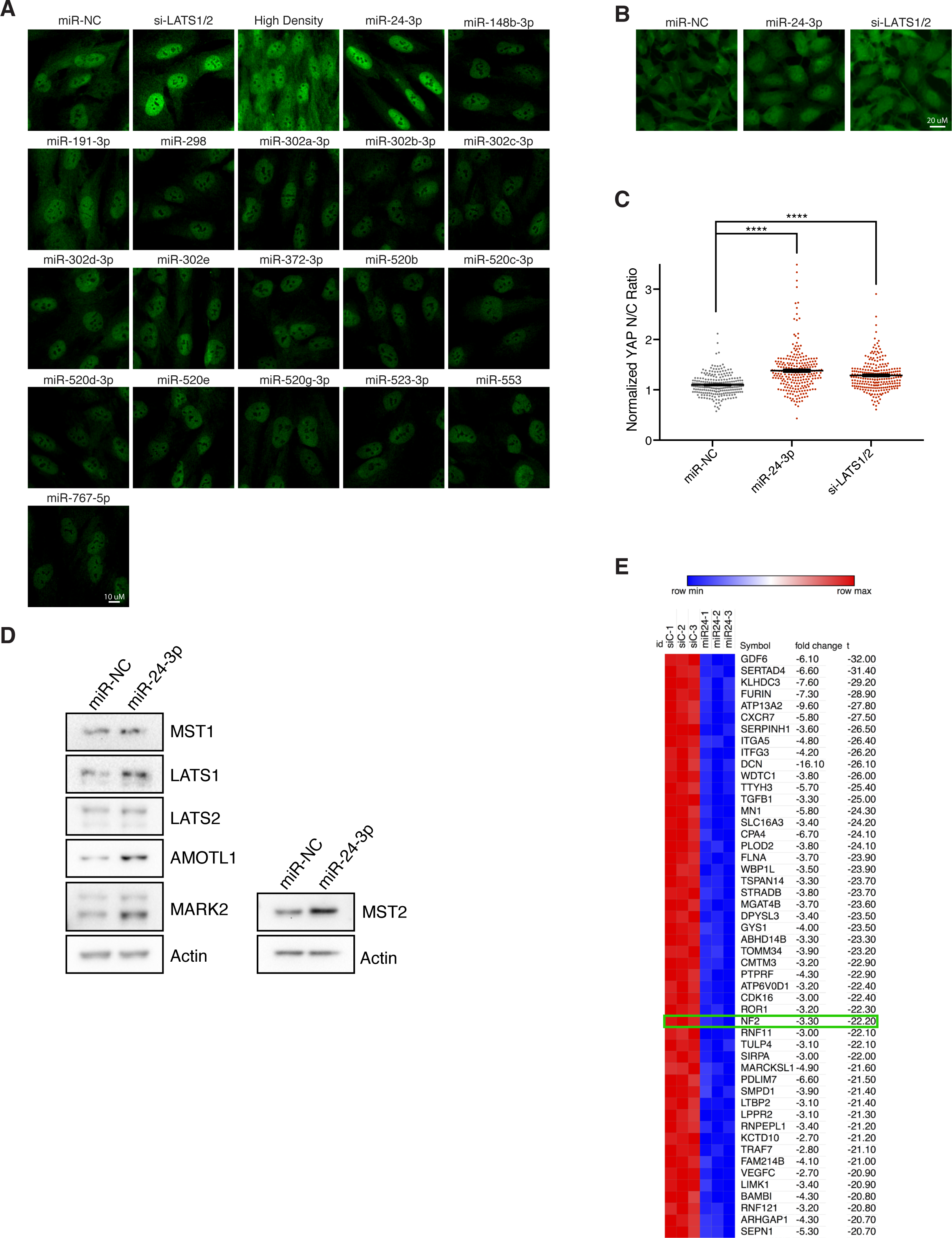
**(A)** Representative fixed immunofluorescence images of YAP (green) in RPE-1 cells transfected with the indicated siRNA/miRNA for 48 hr. **(B)** Representative fixed images of YAP in HEK293A cells transfected with miRNA negative control (miR-NC), miR-24-3p or si-LATS1/2 for 48 hr (n = 3, >200 cells/condition) **(C)** Quantification of the normalized ratio of YAP immunofluorescence intensity in the nucleus:cytoplasm from (B) (3 independent experiments, n > 200 cells/condition, **** p ≤ 0.0001, one-way ANOVA). **(D)** Western blot analysis of indicated proteins in RPE-1 cells transfected with either negative control or miR-24-3p after 48 hrs. **(E)** Heat map of top 50 most downregulated genes, according to t-test, in RPE-1 cells overexpressing miR-24-3p with NF2 highlighted in green. Created using Morpheus matrix visualization software by the Broad Institute (https://software.broadinstitute.org/morpheus/).

## References

Andreassen, P. R., Lohez, O. D., Lacroix, F. B., & Margolis, R. L. (2001). Tetraploid state induces p53-dependent arrest of nontransformed mammalian cells in G1. Mol Biol Cell, 12(5), 1315–1328.

Aylon, Y., Michael, D., Shmueli, A., Yabuta, N., Nojima, H., & Oren, M. (2006). A positive feedback loop between the p53 and Lats2 tumor suppressors prevents tetraploidization. Genes Dev, 20(19), 2687–2700. doi:10.1101/gad.1447006

Bartel, D. P. (2009). MicroRNAs: target recognition and regulatory functions. Cell, 136(2), 215–233. doi:10.1016/j.cell.2009.01.002

Carter, S. B. (1967). Effects of cytochalasins on mammalian cells. Nature, 213(5073), 261–264.

Crockford, A., Zalmas, L. P., Gronroos, E., Dewhurst, S. M., McGranahan, N., Cuomo, M. E.,…Swanton, C. (2017). Cyclin D mediates tolerance of genome-doubling in cancers with functional p53. Ann Oncol, 28(1), 149–156. doi:10.1093/annonc/mdw612

Davoli, T., & de Lange, T. (2011). The causes and consequences of polyploidy in normal development and cancer. Annu Rev Cell Dev Biol, 27, 585–610. doi:10.1146/annurev-cellbio-092910-154234

Davoli, T., & de Lange, T. (2012). Telomere-driven tetraploidization occurs in human cells undergoing crisis and promotes transformation of mouse cells. Cancer Cell, 21(6), 765–776. doi:10.1016/j.ccr.2012.03.044

Dewhurst, S. M., McGranahan, N., Burrell, R. A., Rowan, A. J., Gronroos, E., Endesfelder, D.,…Swanton, C. (2014). Tolerance of whole-genome doubling propagates chromosomal instability and accelerates cancer genome evolution. Cancer Discov, 4(2), 175–185. doi:10.1158/2159-8290.CD-13-0285

Di Leva, G., Piovan, C., Gasparini, P., Ngankeu, A., Taccioli, C., Briskin, D.,…Croce, C. M. (2013). Estrogen mediated-activation of miR-191/425 cluster modulates tumorigenicity of breast cancer cells depending on estrogen receptor status. PLoS Genet, 9(3), e1003311. doi:10.1371/journal.pgen.1003311

Fava, L. L., Schuler, F., Sladky, V., Haschka, M. D., Soratroi, C., Eiterer, L.,…Villunger, A. (2017). The PIDDosome activates p53 in response to supernumerary centrosomes. Genes Dev, 31(1), 34–45. doi:10.1101/gad.289728.116

Fujiwara, T., Bandi, M., Nitta, M., Ivanova, E. V., Bronson, R. T., & Pellman, D. (2005). Cytokinesis failure generating tetraploids promotes tumorigenesis in p53-null cells. Nature, 437(7061), 1043–1047. doi:10.1038/nature04217

Ganem, N. J., Cornils, H., Chiu, S. Y., O’Rourke, K. P., Arnaud, J., Yimlamai, D.,…Pellman, D. (2014). Cytokinesis failure triggers hippo tumor suppressor pathway activation. Cell, 158(4), 833–848. doi:10.1016/j.cell.2014.06.029

Ganem, N. J., Godinho, S. A., & Pellman, D. (2009). A mechanism linking extra centrosomes to chromosomal instability. Nature, 460(7252), 278–282. doi:10.1038/nature08136

Ganem, N. J., & Pellman, D. (2007). Limiting the proliferation of polyploid cells. Cell, 131(3), 437–440. doi:10.1016/j.cell.2007.10.024

Ganem, N. J., Storchova, Z., & Pellman, D. (2007). Tetraploidy, aneuploidy and cancer. Curr Opin Genet Dev, 17(2), 157–162. doi:10.1016/j.gde.2007.02.011

Giglio, S., Cirombella, R., Amodeo, R., Portaro, L., Lavra, L., & Vecchione, A. (2013). MicroRNA miR-24 promotes cell proliferation by targeting the CDKs inhibitors p27Kip1 and p16INK4a. J Cell Physiol, 228(10), 2015–2023. doi:10.1002/jcp.24368

Hatziapostolou, M., Polytarchou, C., Aggelidou, E., Drakaki, A., Poultsides, G. A., Jaeger, S. A.,…Iliopoulos, D. (2011). An HNF4alpha-miRNA inflammatory feedback circuit regulates hepatocellular oncogenesis. Cell, 147(6), 1233–1247. doi:10.1016/j.cell.2011.10.043

Jansson, M. D., & Lund, A. H. (2012). MicroRNA and cancer. Mol Oncol, 6(6), 590–610. doi:10.1016/j.molonc.2012.09.006

Kerimis, D., Kontos, C. K., Christodoulou, S., Papadopoulos, I. N., & Scorilas, A. (2017). Elevated expression of miR-24-3p is a potentially adverse prognostic factor in colorectal adenocarcinoma. Clin Biochem, 50(6), 285–292. doi:10.1016/j.clinbiochem.2016.11.034

Krzywicka-Racka, A., & Sluder, G. (2011). Repeated cleavage failure does not establish centrosome amplification in untransformed human cells. J Cell Biol, 194(2), 199–207. doi:10.1083/jcb.201101073

Kuffer, C., Kuznetsova, A. Y., & Storchova, Z. (2013). Abnormal mitosis triggers p53-dependent cell cycle arrest in human tetraploid cells. Chromosoma, 122(4), 305–318. doi:10.1007/s00412-013-0414-0

Lim, S., & Ganem, N. J. (2014). Tetraploidy and tumor development. Oncotarget, 5(22), 10959–10960. doi:10.18632/oncotarget.2790

Liu, W., Wang, S., Zhou, S., Yang, F., Jiang, W., Zhang, Q., & Wang, L. (2017). A systems biology approach to identify microRNAs contributing to cisplatin resistance in human ovarian cancer cells. Mol Biosyst, 13(11), 2268–2276. doi:10.1039/c7mb00362e

McKinley, K. L., & Cheeseman, I. M. (2017). Large-Scale Analysis of CRISPR/Cas9 Cell-Cycle Knockouts Reveals the Diversity of p53-Dependent Responses to Cell-Cycle Defects. Dev Cell, 40(4), 405–420 e402. doi:10.1016/j.devcel.2017.01.012

Meng, Z., Moroishi, T., & Guan, K. L. (2016). Mechanisms of Hippo pathway regulation. Genes Dev, 30(1), 1–17. doi:10.1101/gad.274027.115

Meng, Z., Moroishi, T., Mottier-Pavie, V., Plouffe, S. W., Hansen, C. G., Hong, A. W.,…Guan, K. L. (2015). MAP4K family kinases act in parallel to MST1/2 to activate LATS1/2 in the Hippo pathway. Nat Commun, 6, 8357. doi:10.1038/ncomms9357

Mootha, V. K., Lindgren, C. M., Eriksson, K. F., Subramanian, A., Sihag, S., Lehar, J.,…Groop, L. C. (2003). PGC-1alpha-responsive genes involved in oxidative phosphorylation are coordinately downregulated in human diabetes. Nat Genet, 34(3), 267–273. doi:10.1038/ng1180

Moroishi, T., Hansen, C. G., & Guan, K. L. (2015). The emerging roles of YAP and TAZ in cancer. Nat Rev Cancer, 15(2), 73–79. doi:10.1038/nrc3876

Moroishi, T., Park, H. W., Qin, B., Chen, Q., Meng, Z., Plouffe, S. W.,…Guan, K. L. (2015). A YAP/TAZ-induced feedback mechanism regulates Hippo pathway homeostasis. Genes Dev, 29(12), 1271–1284. doi:10.1101/gad.262816.115

Nagpal, N., & Kulshreshtha, R. (2014). miR-191: an emerging player in disease biology. Front Genet, 5, 99. doi:10.3389/fgene.2014.00099

Pan, D. (2007). Hippo signaling in organ size control. Genes Dev, 21(8), 886–897. doi:10.1101/gad.1536007

Panopoulos, A., Pacios-Bras, C., Choi, J., Yenjerla, M., Sussman, M. A., Fotedar, R., & Margolis, R. L. (2014). Failure of cell cleavage induces senescence in tetraploid primary cells. Mol Biol Cell, 25(20), 3105–3118. doi:10.1091/mbc.E14-03-0844

Park, Y. Y., Sohn, B. H., Johnson, R. L., Kang, M. H., Kim, S. B., Shim, J. J.,…Lee, J. S. (2016). Yes-associated protein 1 and transcriptional coactivator with PDZ-binding motif activate the mammalian target of rapamycin complex 1 pathway by regulating amino acid transporters in hepatocellular carcinoma. Hepatology, 63(1), 159–172. doi:10.1002/hep.28223

Patel, S. H., Camargo, F. D., & Yimlamai, D. (2017). Hippo Signaling in the Liver Regulates Organ Size, Cell Fate, and Carcinogenesis. Gastroenterology, 152(3), 533–545. doi:10.1053/j.gastro.2016.10.047

Potapova, T. A., Seidel, C. W., Box, A. C., Rancati, G., & Li, R. (2016). Transcriptome analysis of tetraploid cells identifies cyclin D2 as a facilitator of adaptation to genome doubling in the presence of p53. Mol Biol Cell, 27(20), 3065–3084. doi:10.1091/mbc.E16-05-0268

Sakaue-Sawano, A., Kurokawa, H., Morimura, T., Hanyu, A., Hama, H., Osawa, H.,…Miyawaki, A. (2008). Visualizing spatiotemporal dynamics of multicellular cell-cycle progression. Cell, 132(3), 487–498. doi:10.1016/j.cell.2007.12.033

Selmecki, A. M., Maruvka, Y. E., Richmond, P. A., Guillet, M., Shoresh, N., Sorenson, A. L.,…Pellman, D. (2015). Polyploidy can drive rapid adaptation in yeast. Nature, 519(7543), 349–352. doi:10.1038/nature14187

Shenk, E. M., & Ganem, N. J. (2016). Generation and Purification of Tetraploid Cells. Methods Mol Biol, 1413, 393–401. doi:10.1007/978-1-4939-3542-0_24

Silkworth, W. T., Nardi, I. K., Scholl, L. M., & Cimini, D. (2009). Multipolar spindle pole coalescence is a major source of kinetochore mis-attachment and chromosome mis-segregation in cancer cells. PLoS One, 4(8), e6564. doi:10.1371/journal.pone.0006564

Storchova, Z., Breneman, A., Cande, J., Dunn, J., Burbank, K., O’Toole, E., & Pellman, D. (2006). Genome-wide genetic analysis of polyploidy in yeast. Nature, 443(7111), 541–547. doi:10.1038/nature05178

Storchova, Z., & Kuffer, C. (2008). The consequences of tetraploidy and aneuploidy. J Cell Sci, 121(Pt 23), 3859–3866. doi:10.1242/jcs.039537

Subramanian, A., Tamayo, P., Mootha, V. K., Mukherjee, S., Ebert, B. L., Gillette, M. A.,…Mesirov, J. P. (2005). Gene set enrichment analysis: a knowledge-based approach for interpreting genome-wide expression profiles. Proc Natl Acad Sci U S A, 102(43), 15545–15550. doi:10.1073/pnas.0506580102

Sulas, P., Di Tommaso, L., Novello, C., Rizzo, F., Rinaldi, A., Weisz, A.,…Roncalli, M. (2017). A large set of miRNAs is dysregulated since the earliest steps of human hepatocellular carcinoma development. Am J Pathol. doi:10.1016/j.ajpath.2017.10.024

Yu, F. X., Zhao, B., & Guan, K. L. (2015). Hippo Pathway in Organ Size Control, Tissue Homeostasis, and Cancer. Cell, 163(4), 811–828. doi:10.1016/j.cell.2015.10.044

Yuan, Y., Kluiver, J., Koerts, J., de Jong, D., Rutgers, B., Abdul Razak, F. R.,…van den Berg, A. (2017). miR-24-3p Is Overexpressed in Hodgkin Lymphoma and Protects Hodgkin and Reed-Sternberg Cells from Apoptosis. Am J Pathol, 187(6), 1343–1355. doi:10.1016/j.ajpath.2017.02.016

Zack, T. I., Schumacher, S. E., Carter, S. L., Cherniack, A. D., Saksena, G., Tabak, B.,…Beroukhim, R. (2013). Pan-cancer patterns of somatic copy number alteration. Nat Genet, 45(10), 1134–1140. doi:10.1038/ng.2760

Zanconato, F., Forcato, M., Battilana, G., Azzolin, L., Quaranta, E., Bodega, B.,…Piccolo, S. (2015). Genome-wide association between YAP/TAZ/TEAD and AP-1 at enhancers drives oncogenic growth. Nat Cell Biol, 17(9), 1218–1227. doi:10.1038/ncb3216

Zhang, S., Chen, Q., Liu, Q., Li, Y., Sun, X., Hong, L.,…Zhou, D. (2017). Hippo Signaling Suppresses Cell Ploidy and Tumorigenesis through Skp2. Cancer Cell, 31(5), 669–684 e667. doi:10.1016/j.ccell.2017.04.004

Zhou, Y., Frings, O., Branca, R. M., Boekel, J., le Sage, C., Fredlund, E.,…Orre, L. M. (2017). microRNAs with AAGUGC seed motif constitute an integral part of an oncogenic signaling network. Oncogene, 36(6), 731–745. doi:10.1038/onc.2016.242

